# Role of Lateral Inhibition on Visual Number Sense

**DOI:** 10.1101/2021.09.19.460638

**Authors:** Yiwei Zhou, Huanwen Chen, Yijun Wang

## Abstract

Lateral inhibition is a basic principle of information processing and widely exists in the human and animal nervous systems. Lateral inhibition is also involved in processing visual information because it travels through the retina, primary visual cortex, and visual nervous system. This finding suggests that lateral inhibition is associated with visual number sense in humans and animals. Here, we show a number-sensing neural network model based on lateral inhibition. The model can reproduce the size and distance effects of the output response of human and animal number-sensing neurons when the network connection weights are set randomly without adjustment. The number sense of the model disappears when lateral inhibition is removed. Our study shows that the first effect of lateral inhibition is to strengthen the linear correlation between the total response intensity of the input layer and the number of objects. The second one is to allow the output cells to prefer different numbers. Results indicate that lateral inhibition plays an indispensable role in untrained spontaneous number sense.

## INTRODUCTION

Lateral inhibition exists widely in the human and animal nervous systems and is the basis of brain information processing. Lateral inhibition is also present in the primary visual cortex (Lu and Zuo, 2017) and the neocortex (Zhou and Yu, 2018), which are associated with visual processing. The excitatory neurons inhibit the activity of surrounding neurons when the neurons in the two cerebral cortex are stimulated. The intensity of inhibition decays exponentially as the distance between neurons increases, thereby forming a “Mexican hat” interaction relationship, in which the center is excited and the surroundings are inhibited (Field et al., 2020). This interaction can destabilize the attractor state and generate rich responses that encode and store different characteristics of the duration, intensity, and quantity of the external stimulus (Miller, 2013). Previous studies have shown that lateral inhibition plays an important role in the generation of number sense (Sengupta et al., 2014), and other studies have shown that the width of lateral inhibition can affect the encoding mode of number preference neurons (Chen et al., 2020). Transcranial alternating current stimulation of the human parietal lobe (Labree et al., 2020) and quantitative judgment experiments (Clayton and Gilmore, 2015) have also shown that lateral inhibition can affect the number processing ability of organisms. Therefore, lateral inhibition may be a factor of number sense, and studying the role of lateral inhibition in object number information processing is necessary.

At present, many number sense models based on artificial neural networks adopt the function of lateral inhibition directly or indirectly and can reproduce the corresponding experimental results qualitatively or quantitatively (Sengupta et al., 2014; Dehaene and Changeux, 1993; Hannagan et al., 2017; Nasr et al., 2019; Testolin et al., 2020; Nieder, 2016). However, the neural network model cannot systematically explain the role of lateral inhibition in the number sense generation process due to the following reasons. First, most models only use lateral inhibition without analyzing its role (Chen et al., 2020; Dehaene and Changeux, 1993; Hannagan et al., 2017; Nasr et al., 2019; Testolin et al., 2020). Second, humans and animals can estimate the number of objects without visual training, but some models should be trained to adjust the weights between cells to achieve number sense; thus, these models cannot be used to analyze the effect of lateral inhibition on number sense (Nieder, 2016; Burr et al., 2017). Third, some researchers use the deep convolutional neural network computing model with complex structures to solve the problem of number sense in complex real visual scenes (Nasr et al., 2019). Analyzing the effect of lateral inhibition alone is difficult due to the large number of parameters in these models. Therefore, constructing a model based on lateral inhibition and a simple network structure is necessary to study the effect of lateral inhibition without training.

A number sense model based on lateral inhibition is constructed to systematically explain the role of lateral inhibition in number sense generation. The model is a double-layer neural network, which contains only network connection weight and lateral inhibition function, which is beneficial to analyze the role of lateral inhibition. In the case of untrained model (randomly set network connection weights without adjustment), the output response was observed to judge the number sensing ability of the model. On this basis, the role of lateral inhibition in each layer of the neural network is analyzed by canceling the lateral inhibition function.

## METHODS

### Stimulus Set

To test the response of the model to different numbers of objects, the model was tested with three different stimulus sets, as follows: one standard set and two control sets for nondigital visual stimulus cues. In the standard set, the diameter of the circumferential circle of all objects is randomly generated within seven pixels. In the first control set, the total area of objects in each image is 1000 pixels. In the second control set, when the number of objects contained in the image is greater than 4, the convex hull of the object is a regular pentagon with an outer circle of 60 pixels in diameter. Each image in the stimulus set contained 64 × 64 pixels, and the intensity of the stimulus for each pixel ranged from 0 to 1. Each stimulus set has 900 images, containing *n* = 1,2,4,6,8… …28,30 objects, and each number is represented by 30 different images. The shapes of the objects are randomly generated in crosses, rectangles, circles, triangles, squares, and diamonds. Three different stimulus sets are shown in Figure 1A.

**Figure 1.**
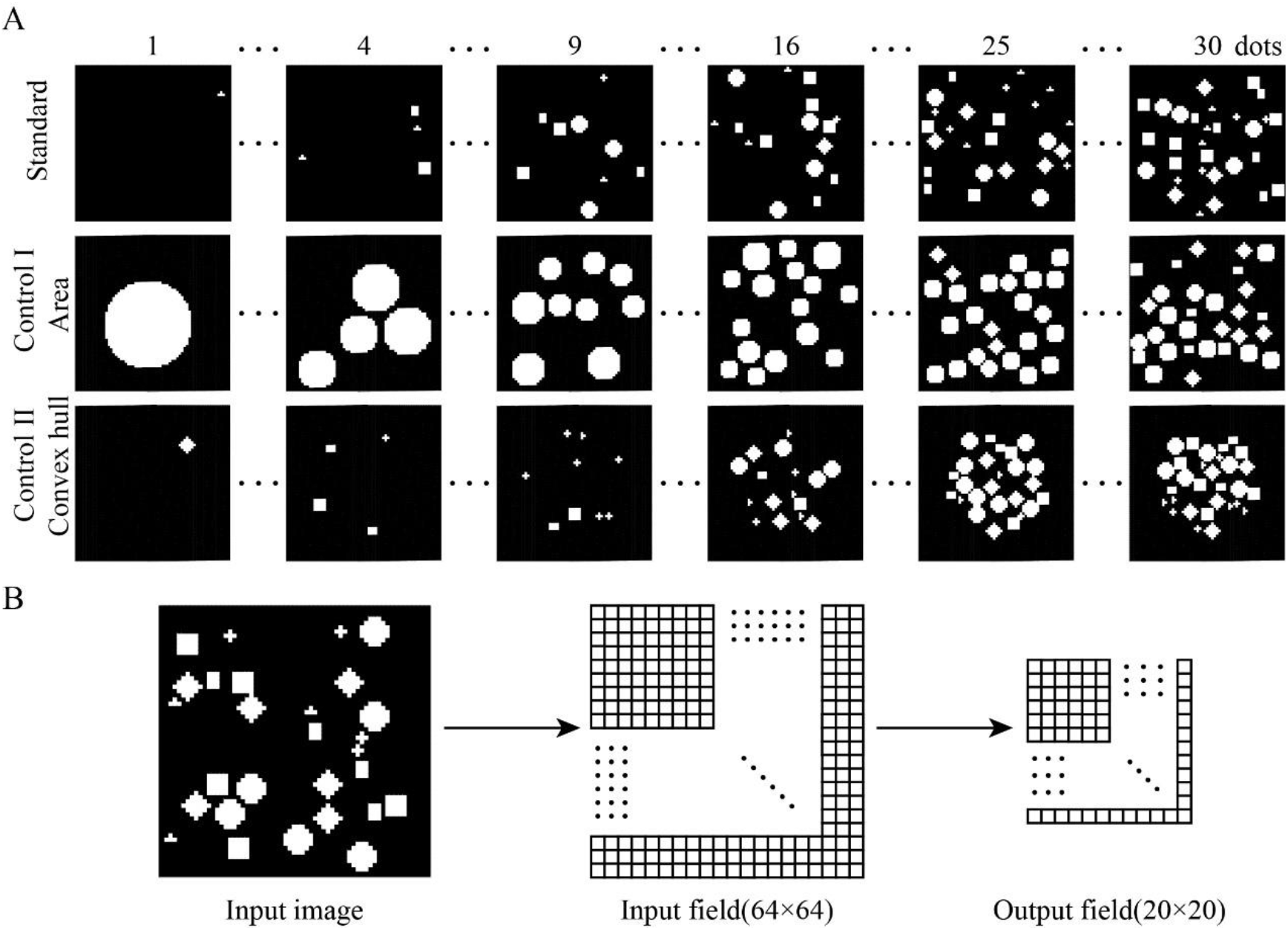
Schematic of stimulus set and neural network structure. (A) Examples of three stimulus sets used to assess the number sense. The diameter of the circumferential circle of the object in the standard set is randomly generated within seven pixels. The stimulus area of each image in control set 1 is 1000 pixels. All objects in control set 2 are inside a regular pentagonal convex hull with a circular diameter of 60 pixels. (B) Number sensing model based on lateral inhibition. The model consists of a double-layer neural network. The input layer has 64 × 64 cells, each cell corresponds to a pixel, and the output layer has 20 × 20 cells. The cells at the same layer inhibit each other, and the cells at different layers are fully connected.

### Model by Lateral Inhibition

On TensorFlow, an open-source machine learning platform, Python programming language, is used to build a number sensing model based on lateral inhibition. The model consists of two layers of neural network (Figure 1B). The first layer of the neural network has 64 × 64 cells, and each cell corresponds to a pixel of the input image. The second layer network has 20 × 20 cells. The cells of the two-layer network are fully connected, and the initial weights follow the Gaussian distribution of *μ* = 0.5, σ^2^ = 0.1. This distribution pattern is qualitatively the same as the synaptic weight distribution observed in biological experiments (Peng et al., 2017).

Each cell is affected by the lateral inhibition of other cells in the same layer network. After the cells of the first-layer neural network receive the visual image input, all cells simultaneously produce lateral inhibition to the surrounding cells. The inhibition intensity is a function of the Euclidean distance *R* between cells and the output intensity *a_x,y_* of cells; it conforms to the characteristics of the Gaussian curve. When the output intensity *a_x,y_* of cells is constant, the inhibition intensity between cells decreases with the increase in *R*.

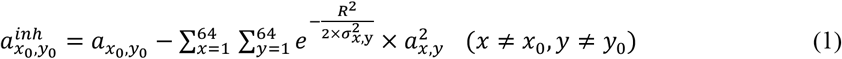

In Formula (1), *a_x_0__,_y_0__* is the input of the network cell of line *x_0_* and column *y_0_* in the first-layer neural network. 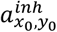 represents the output of the network unit after lateral inhibition. *a_x,y_* is the input of the network cell at row *x* and column *y* in the first-layer neural network. The intensity of lateral inhibition should be higher than that of the excitatory stimulus given that the neural network should avoid overexcitation (Hattori et al., 2017; Scharfman et al., 2014). Therefore, the inhibitory intensity of any cell on the surrounding cells was set to be twice as strong as its input stimulus a_x_,_y_. *R* represents the Euclidean distance between network cells *a_x_0__,_y_0__* and *a_x,y_*. Standard deviation *σ_x,y_* determines the extent of *a_x,y_* lateral inhibition. The lateral inhibition range of each neuron in the cerebral cortex is different (Olson et al., 2021); thus, *σ_x,y_* corresponding to each cell is randomly generated and follows the Gaussian distribution. The expectation and standard deviation of Gaussian distribution were determined by parameter fitting to qualitatively reproduce the experimental results of number sensing neurons in the prefrontal cortex of monkeys (Nieder et al., 2007), and *μ* = 0.67, *σ* = 0.40 were selected.

The output response of each network cell in the first layer neural network is multiplied by the corresponding weight and summed to obtain the input of each network cell in the second layer neural network.

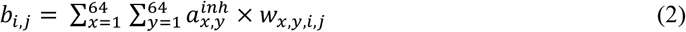

In formula (2), *b_i,j_* is the input of *i* row and *j* column network cell in the second layer neural network. 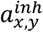 is the output response of the network cell at row *x* and column *y* in the first layer neural network after undergoing lateral inhibition. *w_x,y,i,j_* is the weight between the network cell of row *x* and column *y* in the first layer neural network and the network cell of row *i* and column *j* in the second layer neural network. Each cell of the second layer neural network is simultaneously affected by the lateral inhibition of other cells, and the relationship between inhibition intensity and cell spacing also follows the Gaussian distribution, as follows:

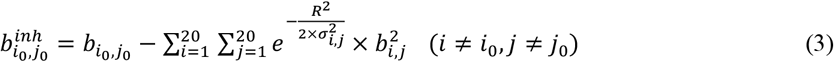

In formula (3), *b_i_0_,j_0__* is the input of the network cell of the row *i_0_* and column *j_0_* in the second layer neural network. 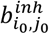 represents the output of the network cell after lateral inhibition. *b_i,j_* is the input of the *i* row *j* column network cell in the second layer neural network. *R* represents the Euclidean distance between network cells *b_i_0__,_j_0__* and *b_i,j_*. The standard deviation *σ_i,j_* corresponding to each cell is randomly generated and follows a Gaussian distribution. The expectation and standard deviation of Gaussian distribution are determined as *μ* = 0.3, *σ* = 0.3 by parameter fitting to reproduce the results of the biological experiment qualitatively (Nieder and Merten, 2007).

### Number Sense Detection

The standard set and the two control sets were input into the neural network to record the response of each cell in the output layer when different images were input. The number of maximum responses is considered the cell preferences, and the number of cells that favor each number is counted. First, to observe whether the response curves in linear scale has scale effect and distance effect, the output responses of the cells with the same quantity preference are averaged to obtain the average response curves with the number of preferences ranging from 1 to 30, and the response curves with the number of preferences *n* = 1,2,4,6,8… …28,30 are drawn in linear scale (*f*(*x*) = x). Second, to study the distribution of the preference number of cells in the output layer, the number of cells with the preference number *n* = 1,2,4,6,8… …28,30 is plotted using a bar graph. Finally, to prove that the output layer cells can better fit Gaussian curves at a nonlinear scale, similar to number sensing neurons (Nieder et al., 2007), Gaussian fitting is performed on the average response curves with a quantity preference of *n* = 1,2,4,6,8… …28,30 on four scales 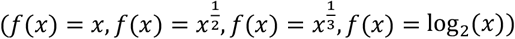. The mean goodness of fit of all the mean response curves at four scales was plotted using a bar graph. At the same time, scatter plots were used to plot the variance of Gaussian fitting of all response curves at four scales, and the relationship between the number of preferences and standard deviation was linearly fitted.

## RESULTS

### The Input Layer and Output Layer Have a Number Sensing Model of Lateral Inhibition

In our neural network model, when both network layers have the function of lateral inhibition, the result of number sense is shown in Figure 2. The left figure, middle figure, and right figure show the number sense results of the model with input of standard set and two control sets. As shown in Figure 2A, the output cell produces a maximum response when the preferred quantity is input and declines in response to other quantities. The larger difference between the number of preferences and other numbers, the larger response gap between the two indicates that it can be easier to distinguish. Therefore, the response curves of the output cell have a distance effect. Figure 2A shows that when the number of objects deviates from the number of preferences, the larger number of preferences of output cells indicates slower decline of output response. Therefore, when the distance between two numbers is fixed, the smaller number indicates that the two numbers are easier to distinguish. The response curve satisfies the size effect.

**Figure 2.**
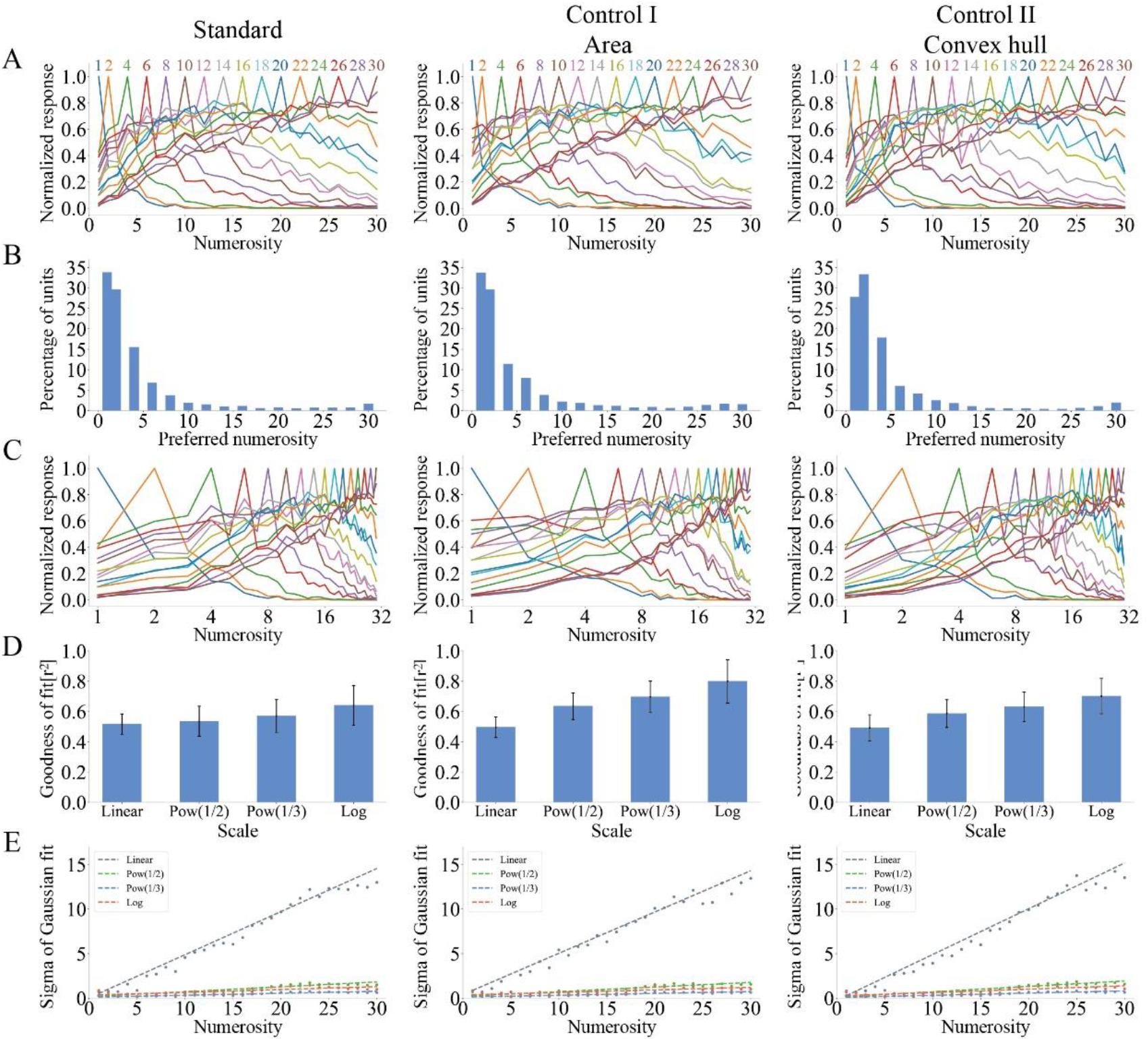
Output response of a model based on lateral inhibition under three different stimulus sets. (A) Average response curves of output layer cells in linear coordinates. The horizontal axis is the number of objects in the image, and the vertical axis is the average response after normalization. (B) Proportion of cells that favor different numbers. The horizontal axis is the number of objects, and the vertical axis is the percentage of all output cells that favor that number. (C) Average response curves of output layer cells in logarithmic coordinates. The horizontal axis is the number of objects in the image, which is plotted on a logarithmic scale of f(x) = log_2_(*x*), and the vertical axis is the average response after normalization. (D) Goodness of fit of Gaussian fitting of response curves at four scales. At the four scales(f(x) = x^0.5^,f(x) = x^0.33^,f(x) = log_2_(x)), the average response curves with the number of preferences ranging from 1 to 30 were combined by Gaussian fitting, and the goodness of fit was calculated. (E) Standard deviation of Gaussian function for best fitting response curves at four scales. The horizontal axis is the number of preferences, and the vertical axis is the standard deviation. Figure 2, from the left figure to the right figure, shows the output response results of the number sense neural network model when the standard set and two control sets are input.

The preference number of each cell is recorded to examine the proportion of cells with different preference numbers in all cells. Figure 2B shows that, under the action of the three stimulus sets, the number of cells with a preference number less than 5 was the largest. As the number of preferences increases, the number of cells that favor that number decreases. Until the number of objects nears 30, the number of cells favoring that number slowly increases. Therefore, the number of cells with different preferences shows the distribution characteristics of high at both ends and low in the middle. This distribution property indicates that our output cells encode numbers with marker lines (Nieder et al., 2007).

The similarity between the response curves and the Gaussian function at different scales should be studied to quantitatively study the distance effect and size effect of the output cell. Average response curves of preference numbers *n* = 1,2,4,6,8… …28,30 were plotted on a logarithmic scale (*f*(*x*) = log_2_(*x*), Figure 2C). The comparison between Figures 2A and 2C shows that the output cell response curves are more symmetrical on a logarithmic scale. Figures 2D and 2E demonstrate these observations with data. Figure 2D shows that the goodness of fit of Gaussian fitting of response curves at linear scale is the lowest, and the average goodness of fit is *r*^2^ = 50.13%. The goodness of fit of Gaussian fitting of response curves increases with the increase in the nonlinearity of abscissa. The mean goodness of fit on a logarithmic scale is *r*^2^ = 71.37°%. Figure 2E shows that the standard deviation increases with the number of preferences on a linear scale. The average slope of the three stimulus sets is *r*_linear_ = 0.49. In other nonlinear scales, the standard deviation is almost unchanged, and the average slope is *r*_non-linear_ = 0.04. These results are consistent with the experimental data in the prefrontal cortex of monkeys (Nieder and Merten, 2007), indicating that the untrained (randomly set and unadjusted network connection weights) lateral inhibition model has number sense ability.

The lateral inhibition model is compared with the model of HDNN (Nasr et al., 2019). The HDNN consists of 15 layers, including eight convolutional layers, six pooling layers, and one output layer. Each layer requires at least three design parameters of mapping feature number, space size, and kernel function size. Therefore, 45 parameters are required from the design of the network structure. In addition, the weight parameters of the network connection and the parameters of the lateral inhibition function should be considered. Therefore, the number of model parameters to be considered is large, and analyzing the effect of lateral inhibition on number sense is difficult. In addition, Nasr et al. ‘s HDNN (Nasr et al., 2019) requires complex pretraining to produce number sense. The selected training set is the ILSVRC2012 ImageNet data set. This dataset contains 1.2 million images, and the training times are 10 epochs (the training data should be presented in full). Therefore, the training time is longer and does not accord with the spontaneity of visual number sense (Nieder, 2016; Burr et al., 2017). Most existing computational models require the quantity-related training of the connection weights between neurons in advance to obtain the function of number sensing (Miller, 2013; Sengupta et al., 2014; Chen and Verguts, 2012; Chen and Chen, 2017). Comparatively, the proposed lateral inhibition model consists of only two layers of neural networks. The model contains only the weights of the network connection and the parameters of lateral inhibition function. More importantly, the lateral inhibition model can reproduce the distance effect, size effect, and marker line coding (Nieder and Merten, 2007) without any learning or training. Compared with traditional convolutional neural networks and deep neural networks (Nasr et al., 2019; Testolin et al., 2020), the lateral inhibition model has usability and simplicity, which is beneficial to analyze the relationship between lateral inhibition and number sense.

### Numerical Sensing Model of Input Layer without Lateral Inhibition

To analyze the effect of input layer lateral inhibition on number sense, in the absence of input layer lateral inhibition, the stimulus scheme in Figure 2 was used to test the model. The responses of input layer cells were plotted under the conditions of removing and retaining input layer lateral inhibition (Figure 3A). The figure shows that when the input layer has no lateral inhibition, the response of cells is equal to the stimulus intensity of corresponding pixels in the image. When the input layer has lateral inhibition, only the cell at the corner of the object has an output response. Therefore, the image stimuli of objects of different sizes and shapes are represented by the response of a few cells after undergoing lateral inhibition. As a result, the positive correlation between the total response strength of the input layer and the number of objects is strengthened.

**Figure 3.**
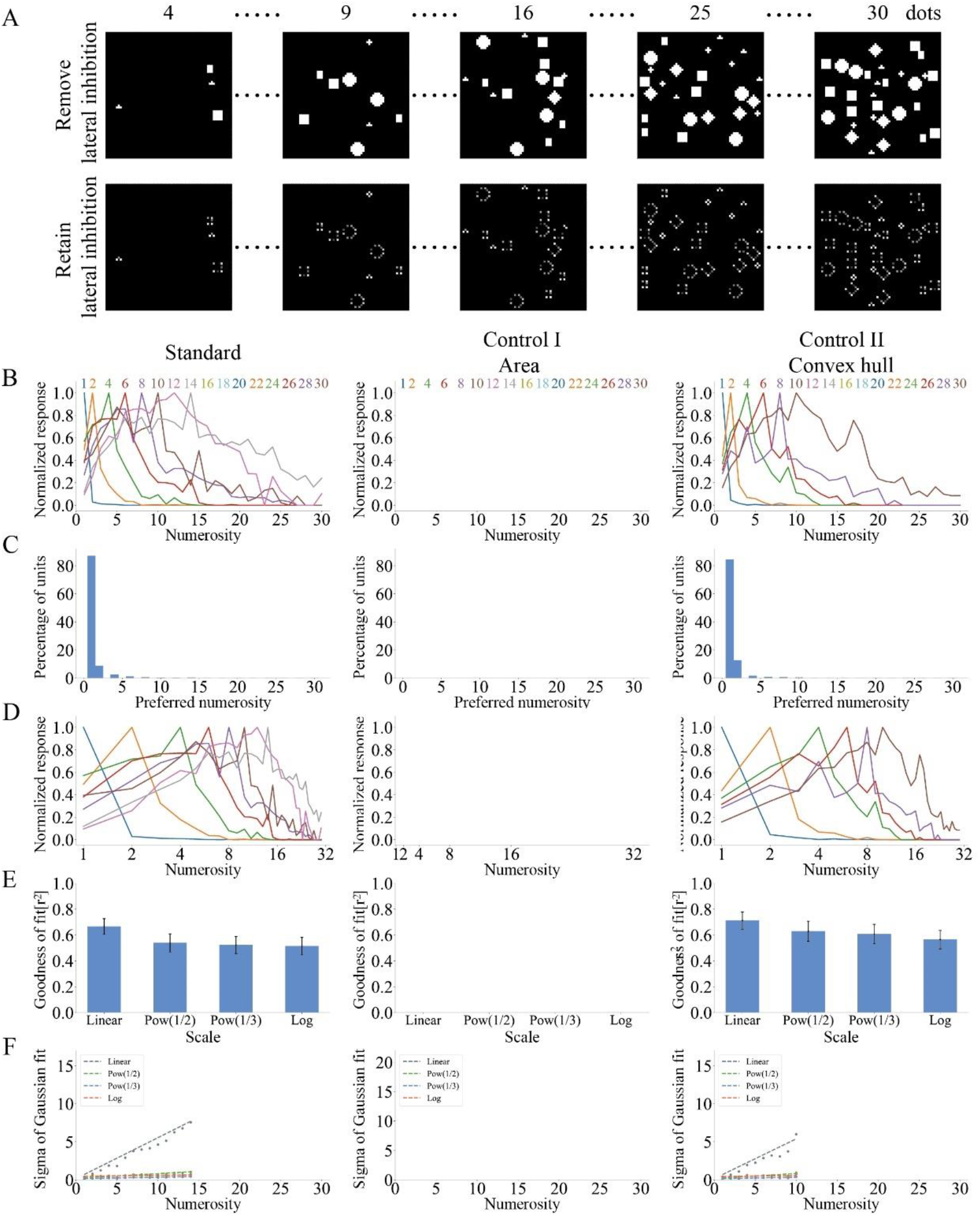
Output response of a neural network model without lateral inhibition at the input layer under three types of stimulus sets. (A) Response of the input layer cell in the two cases of deleting and maintaining the input layer lateral inhibition. The upper layer is the response of the input layer cells after deleting the input layer lateral inhibition. The lower layer is the response of input layer cells when lateral inhibition is retained. (B) Average response curves of output layer cells in linear coordinates. (C) Distribution of the number of cells that prefer different numbers. (D) Average response curves of output layer cells in logarithmic coordinates. (E) Goodness of fit of Gaussian fitting of response curves at four scales. (F) Standard deviation of the best-fit Gaussian function of the response curves under the four scales.

In the case of no lateral inhibition in the input layer, the response curves of cells in the output layer are shown in Figure 3B. The figure shows that under the stimulation of standard set and control set 2 (convex hull), the model can sense the number of objects less than 5, and the average response curve fluctuates greatly. However, when the total area of the object is fixed (control set 1), the model cannot produce a number sense of the object. Figure 3C also shows that for standard set and control set 2, more than 95% of cells prefer numbers within 5. Figure 3D shows the response curve of the output layer cells on a logarithmic scale (*f*(*x*) = log_2_(*x*)). Compared with Figure 3B, the response curve is more asymmetric on a logarithmic scale. The similarity between the response curve and Gaussian curve at different scales was quantitatively studied. First, the goodness of fit of Gaussian fitting at different scales was calculated (Figure 3E). The results indicate that under the action of standard set and control set 2, the goodness of fit at linear scale is higher, *r*^2^_*standard*_ = 66.52% and *r*^2^_*control*2_ = 71.18%, respectively. The goodness of fit decreases with the increase in the nonlinearity of the abscissa. Second, the relationship between the standard deviation of the best-fitting Gaussian function and the number of preferences at four scales was calculated (Figure 3F). The results show that the standard deviation increases with the increase in preference number in linear scale but remains unchanged in other nonlinear scales.

Figure 2 shows the lack of lateral inhibition in the input layer results in two changes. First, it causes the response curve to fluctuate wildly because the linear correlation coefficient between the total response strength of the input layer and the number of objects decreases when the lateral inhibition of the input layer is removed. The second is the variation in the range of preferred numbers of the cell. For standard set and control set 2, the number of preferences is within 5 because, in the absence of input layer lateral inhibition, the total response strength of the input layer increases, as well as the stimulus intensity of the input layer to the output layer cells. At the same time, the lateral inhibition intensity of output layer cells to other cells increases. Only cells at the edge of the image can produce the maximum response to more than five objects. No number sense can be generated with input from the control set 1 because the total response strength of the input layer is constant when the total area of the object is constant. As a result, the input stimulus intensity of cells in the output layer is unchanged, and the inhibition intensity of each cell to other cells is also unchanged. Therefore, the response of the cells in the output layer is the same when the number of inputs is different, and the number of objects cannot be sensed. The model without inhibition of the input layer did not possess number sense ability because the model could not generate number sense stably when controlling the input of nondigital visual stimulus cues.

### Number Sensing Model for Output Layer without Lateral Inhibition

Three types of stimulus sets were used to test the model without output layer lateral inhibition to study the effect of output layer lateral inhibition on the number sensing ability of the model. The average response curve at a linear scale is shown in Figure 4A. The figure shows that the output response of the model rises with the increase in the number of objects because with the increase in the number of objects, the total response strength of the input layer increases, eventually increasing the response of the cells in the output layer. The distribution of cell numbers with different preferences (Figure 4B) was plotted. The figure shows that some cells also have the maximum response when the number of objects is close to 30 because, after the lateral inhibition of the input layer, the total response intensity of the input layer has an incompletely positive linear correlation with the number of objects. Combined with the influence of the initial weight conforming to the Gaussian distribution, the cell may also have the maximum response to the number of objects less than 30. The average response curve on a logarithmic scale (*f*(*x*) = log_2_ (*x*)) is shown in Figure 4C. The response curves are fitted with Gaussian on linear and nonlinear scales. The figure shows that the goodness of fit of the response curve decreases with the increase in the nonlinear scale of the abscissa (Figure 4D). The linear relationship between the standard deviation of the best-fitting Gaussian function and the number of preferences at different scales (Figure 4E) is qualitatively the same as Figure 2. Compared with Figure 2, the lateral inhibition of the output layer causes the cells of the output layer to have preferences for different numbers. The model without lateral inhibition of the output layer has no number sense ability.

**Figure 4.**
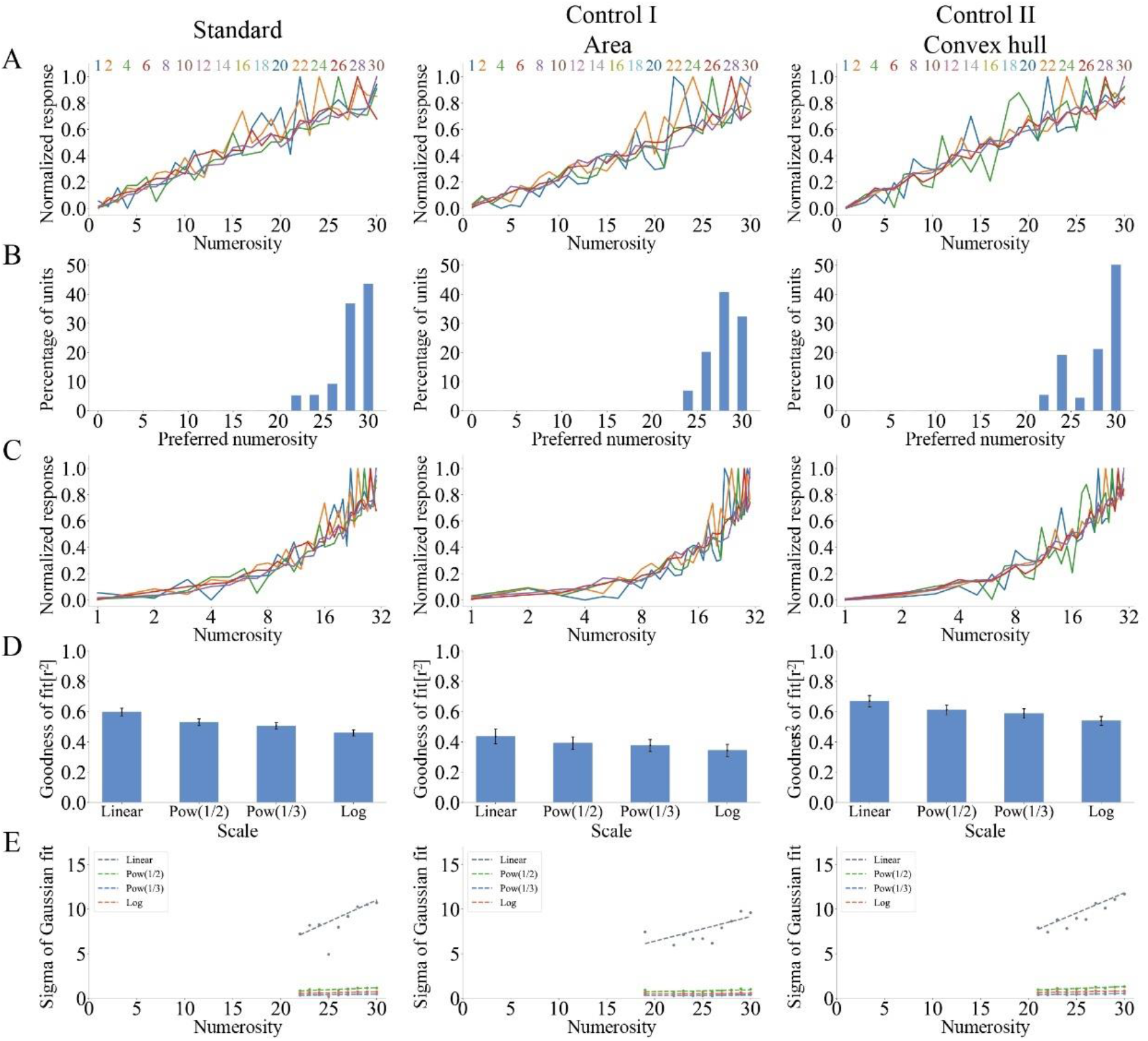
Output response of a neural network model without lateral inhibition at the output layer under three types of stimulus sets. (A) Average response curves of output layer cells in linear coordinates. (B) Distribution of the number of cells that prefer different numbers. (C) Average response curves of output layer cells in logarithmic coordinates. (D) Goodness of fit of Gaussian fitting of response curves at four scales. (E) Standard deviation of Gaussian fitting of response curves at four scales.

### Number Sensing Model without Lateral Inhibition in Input and Output Layers

To investigate whether the model without lateral inhibition has number sensing capability, the lateral inhibition function of the input layer and output layer is removed. The model under the same stimulus as Figure 2 is tested. The average response curves of the model are shown in Figure 5A. The figure shows that the average output response of the model increases with the increase in the number of objects and fluctuates greatly. The network cells of all output layers only respond at a maximum when the number of preferences approaches 30 (Figure 5B). The same finding is true for the average response curve plotted on a logarithmic scale (Figure 5C). Gaussian fitting was performed on the response curves at four scales. Only when control set 1 is input, the goodness of fit increases with the increase in the nonlinearity of the abscissa (Figure 5D). Only on a linear scale does the standard deviation of the best-fitting Gaussian function increases as the number of preferences increases (Figure 5E). The results are consistent with those of the output layer without lateral inhibition. The model is proven to have no number sense when the input layer and output layer lack lateral inhibition. Lateral inhibition is one of the necessary factors to produce number sense.

**Figure 5.**
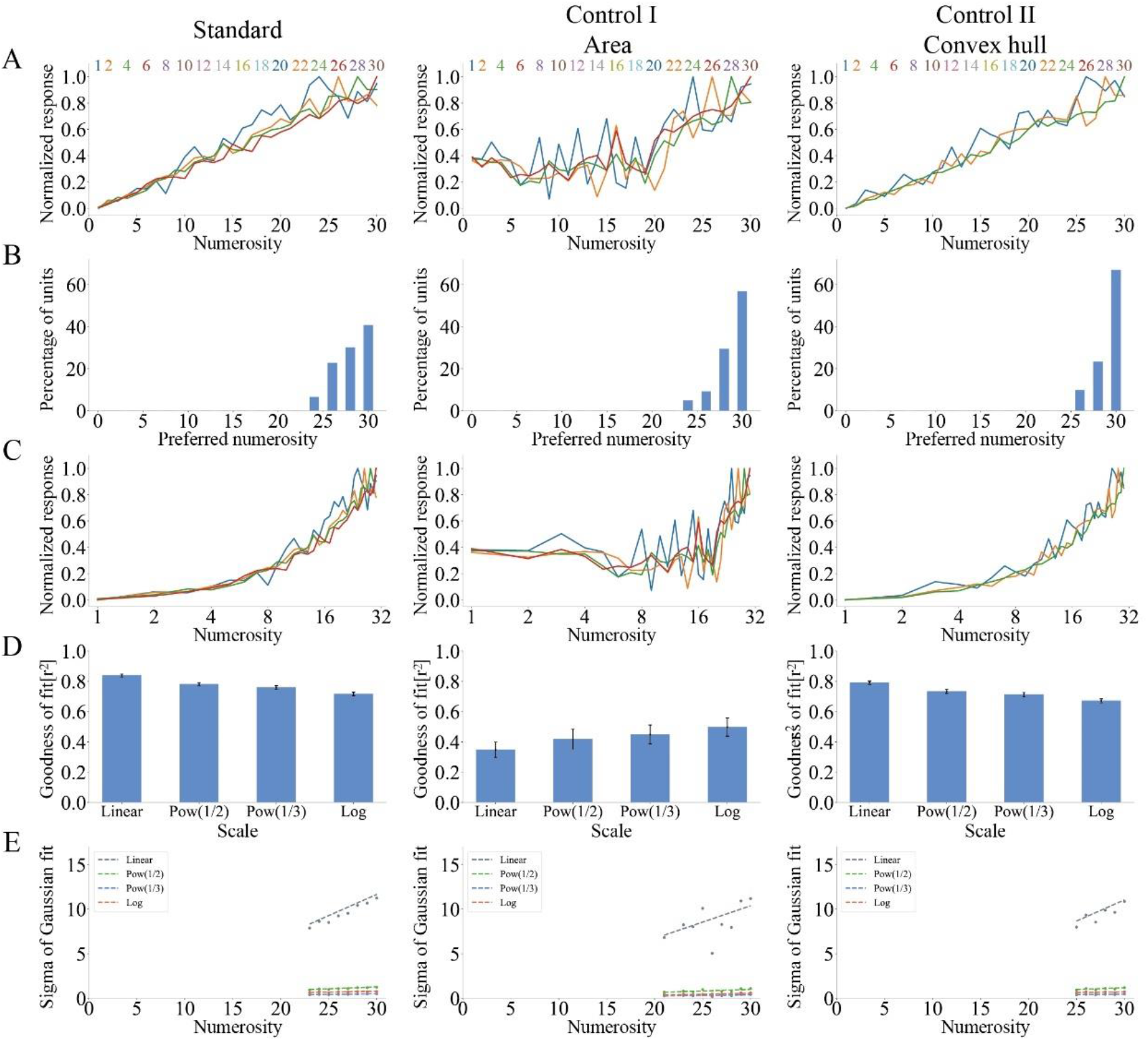
Output response of a neural network model without lateral inhibition in the input–output layer under three stimulus sets. (A) Average response curves of output layer cells in linear coordinates. (B) Distribution of the number of cells that prefer different numbers. (C) Average response curves of output layer cells in logarithmic coordinates. (D) Goodness of fit of Gaussian fitting of response curves at four scales. (E) Standard deviation of Gaussian fitting of response curves at four scales.

## DISCUSSION

Compared with other digital processing network models, our number sensing neural network model has three main advantages. The first point is that we simplify the number of network layers and the number of parameters. Most number sense models use complex deep neural networks and convolutional neural networks to process complex visual images (Nasr et al., 2019; Testolin et al., 2020). On the contrary, our model uses a two-layer neural network structure. The model contains only two parameters, synaptic weight and lateral inhibition, which are helpful to analyze the process of number sense. The second advantage is the spontaneity of the number sense generation process. Different from previous number-sensing models, which relied on quantity-dependent training (Miller, 2013; Sengupta et al., 2014; Chen and Verguts, 2012; Chen and Verguts, 2017), our model does not require pretraining. When the network connection weights are set randomly and no adjustment is made, the model can reproduce the distance effect, size effect, and marker line coding (Nieder and Merten, 2007). Finally, our model mimics the human and animal cerebral cortex and introduces a biophysical lateral inhibition function.

Although lateral inhibition function is adopted in most models (Nasr et al., 2019; Testolin et al., 2020), the specific role of lateral inhibition is not explained. We find that lateral inhibition has different effects in each layer of the network by selectively removing lateral inhibition in the input layer and output layer. The function of the input layer is to improve the correlation between the total stimulus intensity in the input layer and the number of objects because different object images can be represented by the output response of a few cells in the input layer under the effect of lateral inhibition. The role of lateral inhibition in the output layer is to allow the output layer cells to have a preference for different numbers because the output response of cells in the output layer is related not only to excitatory stimuli in the input layer but also to lateral inhibition in surrounding cells. The lateral inhibition intensity increased faster than the excitatory stimulus intensity with the increase in the number of objects. The maximum cell response occurs when the lateral inhibition intensity of the cell is equal to the excitatory stimulus. At the same time, the cells closer to the edge of the neural network suffer from less lateral inhibition. Therefore, as the Euclidean distance between cells and the geometric center of the neural network increases, the number of cell preferences increases. Lateral inhibition is the key to spontaneous number sense.

A customized grayscale image is used as the stimulus set to simplify the number of network layers and parameters. Although the nondigital visual stimuli were controlled, a gap between the stimulus set and the actual visual images still exists. Objects in actual visual images have complex shapes and rich colors. A partial overlap of objects also exists. Therefore, testing the number sensing ability of the neural network model based on lateral inhibition in actual visual scenes is necessary. At the same time, our study shows that a simple neural network with the function of lateral inhibition can achieve spontaneous number sense without training. However, whether the number sense ability can be greatly improved if the neural network is studied and trained is unknown. In addition, the stimulus set contains only a certain number of objects, whereas humans can recognize abstract numbers. Therefore, whether our model can be trained to recognize abstract numbers needs further exploration.

## Author Contributions

Yiwei Zhou: Conceptualization; Data curation; Formal analysis; Investigation; Methodology; Writing— Original draft. Huanwen Chen: Conceptualization; Writing— Review & editing; Yijun Wang: Writing— Review & editing.

## Funding Information

This work was supported by the Hunan Provincial Natural Science Foundation of China (2021JJ30863).

## Notes

### Competing Interest Statement

The authors have declared no competing interest.

